## Supplementary material for "Linalool’s Multifaceted Antimicrobial Potential: Unveiling its Antimicrobial Efficacy and Immunomodulatory Role Against *Saprolegnia parasitica*": Figure S1, Figure S2, and Table S1

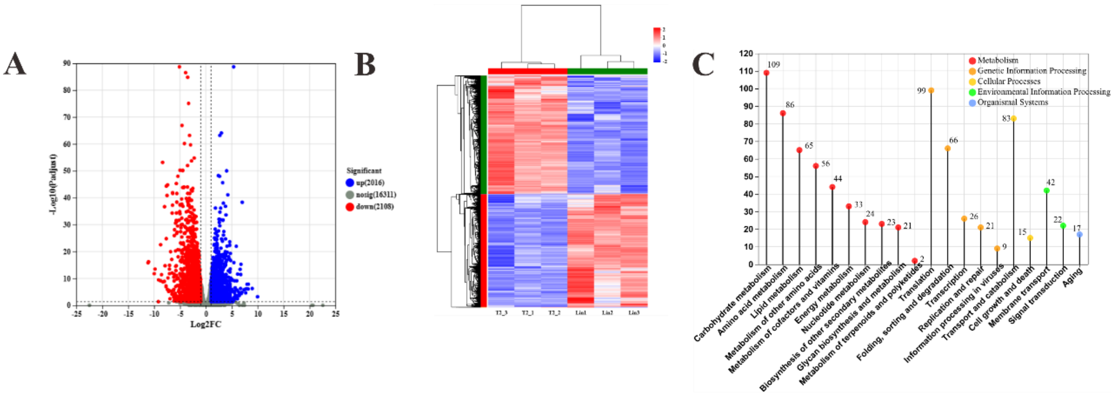


**Figure S1.** Global transcriptomic analysis between the treated (mycelium treated with linalool) and control group (untreated with linalool). A) The number of DEGs. B) Heat map of DEGs. The red cluster represented up regulated genes, whereas the blue cluster represented down regulated genes; C) The KEGG classification of DEGs.

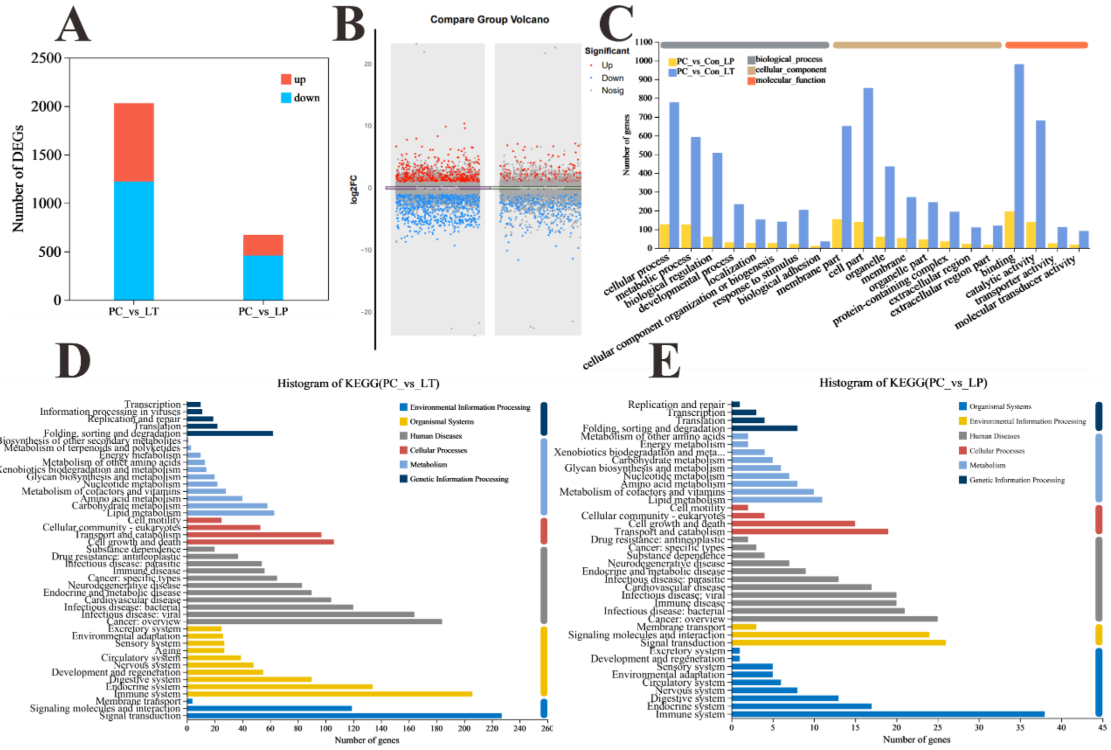


**Figure S2.** Global transcriptomic analysis after *S. parasitica* infection and linalool treatment in grass carp. A) The number of DEGs. B) The scatter diagram of DEGs. C) GO classification of DEGs in each group. D) KEGG classification of DEGs in group LT. E) KEGG classification of DEGs in group LP.

Table S1. Binding energy of linalool with different ribosome proteins.

| Proteins | Binding energy with linaloool |
| --- | --- |
| NOP1 | -6.2kcal/mol |
| DCK1 | -5.0kcal/mol |
| SNU13 | -4.8kcal/mol |
